# Socioeconomic inequality in recent adverse mortality trends in Scotland

**DOI:** 10.1101/542472

**Authors:** Lynda Fenton, Grant Wyper, Gerry McCartney, Jon Minton

## Abstract

**Background:** Gains in life expectancies have stalled in Scotland, as in several other countries, since around 2012. The relationship between stalling mortality improvements and socioeconomic inequalities in health is unclear.

**Methods:** We calculate the percentage improvement in age-standardised mortality rates (ASMR) in Scotland overall, by sex, and by Scottish Index of Multiple Deprivation (SIMD) quintile and gender, for two periods: 2006-2011 and 2012-2017. We then calculate the socioeconomic gradient in improvements for both periods.

**Results:** Between 2006 and 2011, ASMRs fell by 10.6% (10.1% in females; 11.8% in males), but between 2012 and 2017 ASMRs only fell by 2.6% (3.5% in females; 2.0% in males). The socioeconomic gradient in ASMR improvement more than quadrupled, from 0.4% per quintile in 2006-2011 (0.7% in females; 0.6% in males) to 1.7% (2.0% in females; 1.4% in males). Within the most deprived quintile, ASMRs fell in the 2006-2011 period (8.6% overall; 7.2% in females; 9.8% in males), but rose in the 2012-2017 period (by 1.5% overall; 0.7% in females; 2.1% in males).

**Conclusion:** As mortality improvements in Scotland stalled in 2012-2017, socioeconomic gradients in mortality became steeper, with increased mortality rates over this period in the most socioeconomically deprived fifth of the population.

What we already know

- Improvements in mortality rates slowed markedly around 2012 in Scotland and a number of other high-income countries.
- Scotland has large socioeconomic health inequalities, and the absolute gap in premature mortality between most and least deprived has increased since 2013.
- The relationship between stalling mortality improvements and socioeconomic inequalities in health is unclear.

What this study adds

- Stalling in mortality improvement has occurred across the whole population of Scotland, but is most acute in the most socioeconomically deprived areas.
- Mortality improvements went into reverse (i.e. deteriorated) in the most deprived fifth of areas between 2012 and 2017.
- Research to further characterise and explain recent aggregate trends should incorporate consideration of the importance of socioeconomic inequalities within proposed explanations.

## Introduction

Overall mortality trends in Scotland have worsened in recent years, as in a number of other high-income countries.^1–3^ The most recent life expectancy data published (2015-2017) showed that life expectancy fell by 0.1 years for both men and women, compared with the previous period.^4^ Scotland has the lowest level of life expectancy in Western Europe, and absolute inequalities in premature mortality have increased since 2013.^5,6^

High levels of inequality in mortality, especially in the working-age population, have been suggested as an explanation for why overall life expectancy in Scotland has historically lagged behind comparable countries.^7^ The degree to which the recent stalling improvement in mortality rates is associated with changes in health inequalities is unclear.

Age-standardised mortality rates (ASMRs) permit comparison of mortality risks across time and place, in a way that controls for population structure. National Records of Scotland (NRS) publish ASMRs by population quintile, as measured by Scottish Index of Multiple Deprivation (SIMD).^8,9^ We use these recently published data to explore whether trends in mortality inequality have changed in Scotland in a recent six-year period of stalling life expectancies (from 2012 to 2017) compared with the preceding six-years (2006 to 2011).

## Methods

Published ASMRs (using the European standard population 2013) for Scotland, by SIMD^*^ quintile, for males and females of all ages, were downloaded from the NRS website.^8^ The trends in ASMRs from 2001 to 2017 inclusive, by sex, overall and by SIMD quintile, were plotted. Recent analysis of mortality trends in England identified a breakpoint in the early 2010s, and our replication of this analysis for Scottish data identified a contemporaneous breakpoint.^10^ The percentage change in ASMRs from 2006 to 2011 (period 1), and from 2012 to 2017 (period 2), by sex, overall for Scotland, and by SIMD quintile, was calculated. To assess inequalities in trends we regressed the percentage ASMR improvement against SIMD quintile using linear regression for both periods. Each model fit was assessed with the R-squared statistic, and the intercept (predicted percentage improvements in most deprived quintile) and gradient (average increase in percentage improvement associated with move up one quintile category) was calculated.

## Results

Figure 1A shows how ASMRs have changed from 2001 to 2017. In 2006-2011 ASMR improved overall (−10.6%), for females (−10.1%) and males (−11.8%). In 2012-2017 ASMRs improved much more slowly (−2.6% overall; −3.5% in females; −2.0% in males). Between 2001 and 2017, ASMRs fell from 1415 to 1143 per 100,000 overall (1195 to 997 per 100,000 for females; 1735 to 1329 per 100,000 for males), and in all deprivation quintiles.

**Figure 1.**
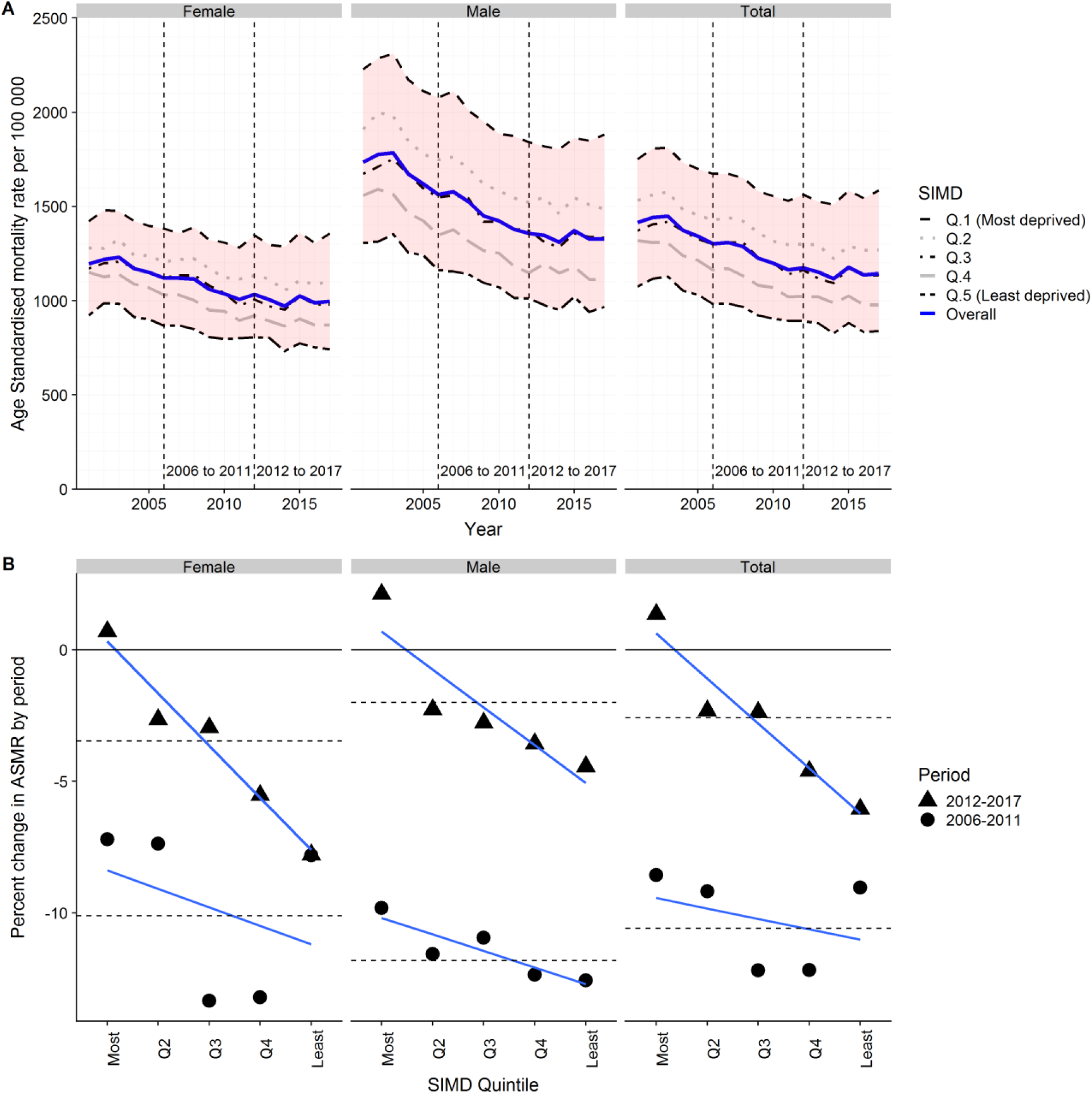
(A) Age-standardised mortality rates, Scotland, 2001-2017, by gender and SIMD quintile. (B) Percentage change in ASMR by period (2006-2011; 2012-2017) by gender and SIMD quintile. Horizontal dashed lines indicate average change in Scotland by period and gender.

ASMR improvements were smaller in the later than earlier period in all deprivation quintiles, except the least deprived females (see Figure 1B and Table 1), and in the latter period worsened (i.e. rose rather than fell) in the most deprived quintile. In both periods, the most deprived SIMD quintile (1) saw the smallest mortality improvements. Figure 1B shows the nature of the relationship between percentage change in mortality rate and SIMD quintile, for the two periods. This shows that there is a steeper socioeconomic gradient in the improvement in mortality in period 2 than in the preceding six years. Among males there is a gradient across quintiles in both periods, but among females the relationship in the period1 appears U-shaped, with the most and least deprived groups having smaller mortality improvements than quintiles 2 and 3.

**Table 1:**
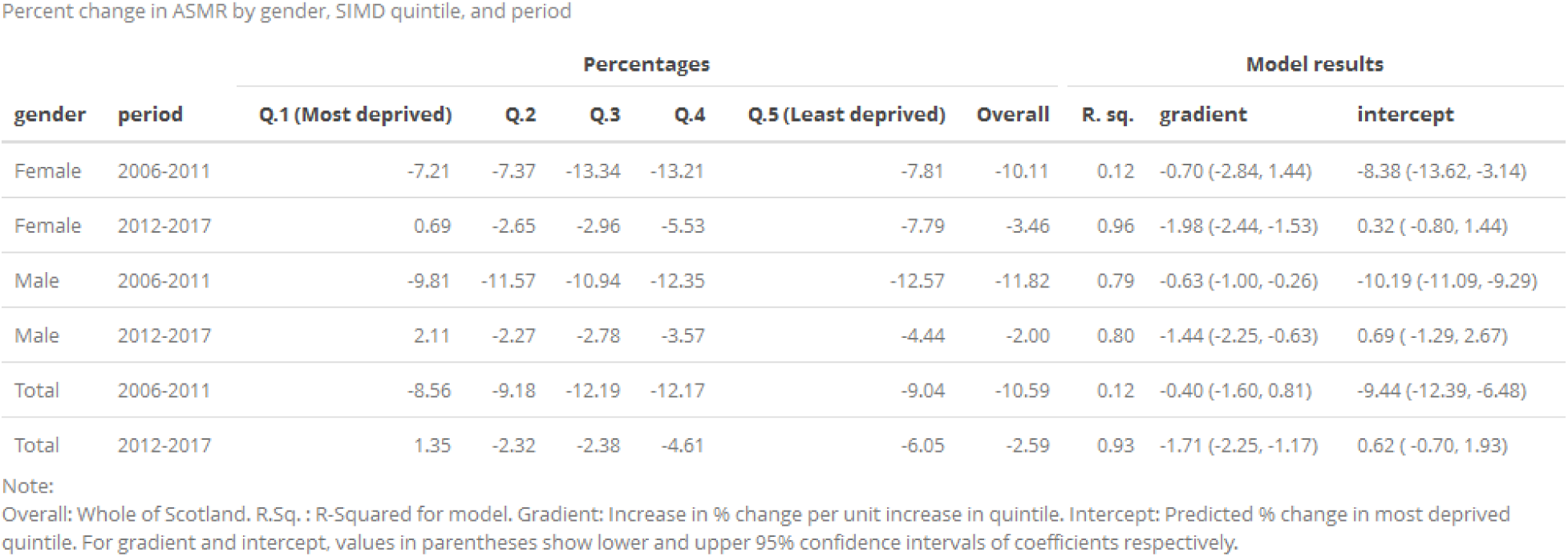
Percentage change in ASMR by gender, SIMD quintile (Q), time period, and results of linear model.

| gender | period | Percentages |  |  |  |  |  | Model results |  |  |
| --- | --- | --- | --- | --- | --- | --- | --- | --- | --- | --- |
|  |  | Q.1 (Most deprived) | Q.2 | Q.3 | Q.4 | Q.5 (Least deprived) | Overall | R. sq. | gradient | intercept |
| Female | 2006-2011 | -7.21 | -7.37 | -13.34 | -13.21 | -7.81 | -10.11 | 0.12 | -0.70 (-2.84, 1.44) | -8.38 (-13.62, -3.14) |
| Female | 2012-2017 | 0.69 | -2.65 | -2.96 | -5.53 | -7.79 | -3.46 | 0.96 | -1.98 (-2.44, -1.53) | 0.32 (-0.80, 1.44) |
| Male | 2006-2011 | -9.81 | -11.57 | -10.94 | -12.35 | -12.57 | -11.82 | 0.79 | -0.63 (-1.00, -0.26) | -10.19 (-11.09, -9.29) |
| Male | 2012-2017 | 2.11 | -2.27 | -2.78 | -3.57 | -4.44 | -2.00 | 0.80 | -1.44 (-2.25, -0.63) | 0.69 (-1.29, 2.67) |
| Total | 2006-2011 | -8.56 | -9.18 | -12.19 | -12.17 | -9.04 | -10.59 | 0.12 | -0.40 (-1.60, 0.81) | -9.44 (-12.39, -6.48) |
| Total | 2012-2017 | 1.35 | -2.32 | -2.38 | -4.61 | -6.05 | -2.59 | 0.93 | -1.71 (-2.25, -1.17) | 0.62 (-0.70, 1.93) |

Note:
Overall: Whole of Scotland. R.Sq.: R-Squared for model. Gradient: Increase in % change per unit increase in quintile. Intercept: Predicted % change in most deprived quintile. For gradient and intercept, values in parentheses show lower and upper 95% confidence intervals of coefficients respectively.

The low R-Squared value (0.12) in the 2006-2011 period, confirms that a linear model is not a good fit for this period, but is a very good fit (0.93) in the 2012-2017 period. The poor fit in the earlier period, and the improved later fit, is largely driven by the gradient in females (R-Squared 0.12 to 0.96), with the fit in males similar in both periods (0.79 to 0.80). The gradient of the fit, which can be considered a measure of the importance of socioeconomic gradient to health inequalities, has increased markedly, from 0.4% per quintile in the 2006-2011 period to 1.7% per quintile in the 2012-2017 period; a ratio of gradients of 4.25, and difference in gradients of 1.3%. The change in ratio of gradients was steeper for females (16.5 compared with 2.3 for males), and the difference in gradients was greater for females (1.3%) than for males (0.8%).

The column marked ‘intercept’ (table 1) is the percentage improvement in quintile 1 (the most deprived quintile) predicted by the linear models. This can be compared with the observed quintile 1 values in the third column of table 1 (and the corresponding points in Figure 1B). In both periods the model predicts greater improvement for this quintile than that which was observed.

## Discussion

This paper shows that in 2012-2017 ASMR improvement in Scotland was much more modest than in the previous six-year period, that ASMR rates worsened in the most deprived fifth of areas, and that the rate of improvement also slowed markedly in the least deprived areas. The socioeconomic gradient in recent mortality rate change is steepest amongst women.

Our findings are similar to those identified in England, which found that the life expectancy gap between the most and least deprived deciles increased between 2001 and 2016, that life expectancy fell for females in the most deprived fifth of areas, and a report by Public Health England showing a widening absolute life expectancy gap by IMD in 2014-16.^11,12^

This paper analysed differences in ASMRs by SIMD quintile. Further research should be conducted to test how dependent our findings are on the choice of health outcome and socioeconomic variable. We used published mortality data which ranks the population by the SIMD. This deprivation index includes a health domain, raising the possibility of circularity in the analysis; however the overall SIMD index is very highly correlated with the income-employment domains which might be used to address this concern. Area-based deprivation analyses misclassify a large proportion of people who are individually deprived and therefore tends to estimate lower inequalities than would be the case with individual measures.

The comparison periods adopted are likely to affect the results, though the use of 2012-2017 for the latter period is justified by analyses demonstrating a change in mortality trends around 2012. The analyses compare the first and last year within each of the two periods, which might not be appropriate if either first or last year in each period are atypical. For this reason a sensitivity analysis comparing values modelled on the trend (rather than observed at the start/end of each period) was conducted, and produced qualitatively similar conclusions (see appendix).

Further research is needed to investigate the drivers of recent adverse mortality trends, and the interaction between mortality trend stalling and health inequalities. Research to further characterise and explain recent aggregate trends should incorporate consideration of the importance of socioeconomic inequalities within proposed explanations.

We have shown that stalling in mortality improvements has occurred across the whole population of Scotland, but is most acute in the most socioeconomically deprived areas. The identification of this important inequalities aspect to recent mortality trends highlights the need for policy-makers to develop responses which seek to undo the fundamental causes of inequality, and which are also proportionate to need, in terms of access to key public services for care and prevention.

## Appendix Sensitivity Analysis

### Introduction

The results presented above compare the first and last year in two six year periods,

2006-2011, and 2012-2017. This approach might lead to unrepresentative results if the ASMR values in the first or last year of either period are unusual or uncharacteristic of the period as a whole. In order to assess whether this is an important issue, the values shown in figure 1B and table 1 were replicated using an alternative method. The alternative method, and results, are shown in this appendix.

### Method

For the alternative method, linear regression was used to regress ASMR against year for males and females overall, and in each deprivation quintile, for both the 2006-2011 period, and the 2012-2017 period. This meant a total of 24 regression models (two genders, five SIMD quintiles plus overall, and two time periods) were produced, each regressing a group’s ASMRs against time for six years. For each group, the fitted values for ASMRs in the first and last year of each period were calculated, and used in place of the observed values in the analyses described in the main report. The results using the fitted values instead of observed values are shown below.

### Results

Figure 2 shows the percentage change in ASMR by period, gender and SIMD quintile, using the within-period fitted values rather than the observed values; horizontal dashed lines indicate the overall percentage change within the period. The results shown in the figure are similar to that produced using the main method, though more clearly suggest that a linear socioeconomic gradient in ASMR improvement was established only in the 2012-2017 period, with a broadly flat (males) or U-shaped (females, total) relationship observed in the 2005-2011 period.

**Figure 2.**
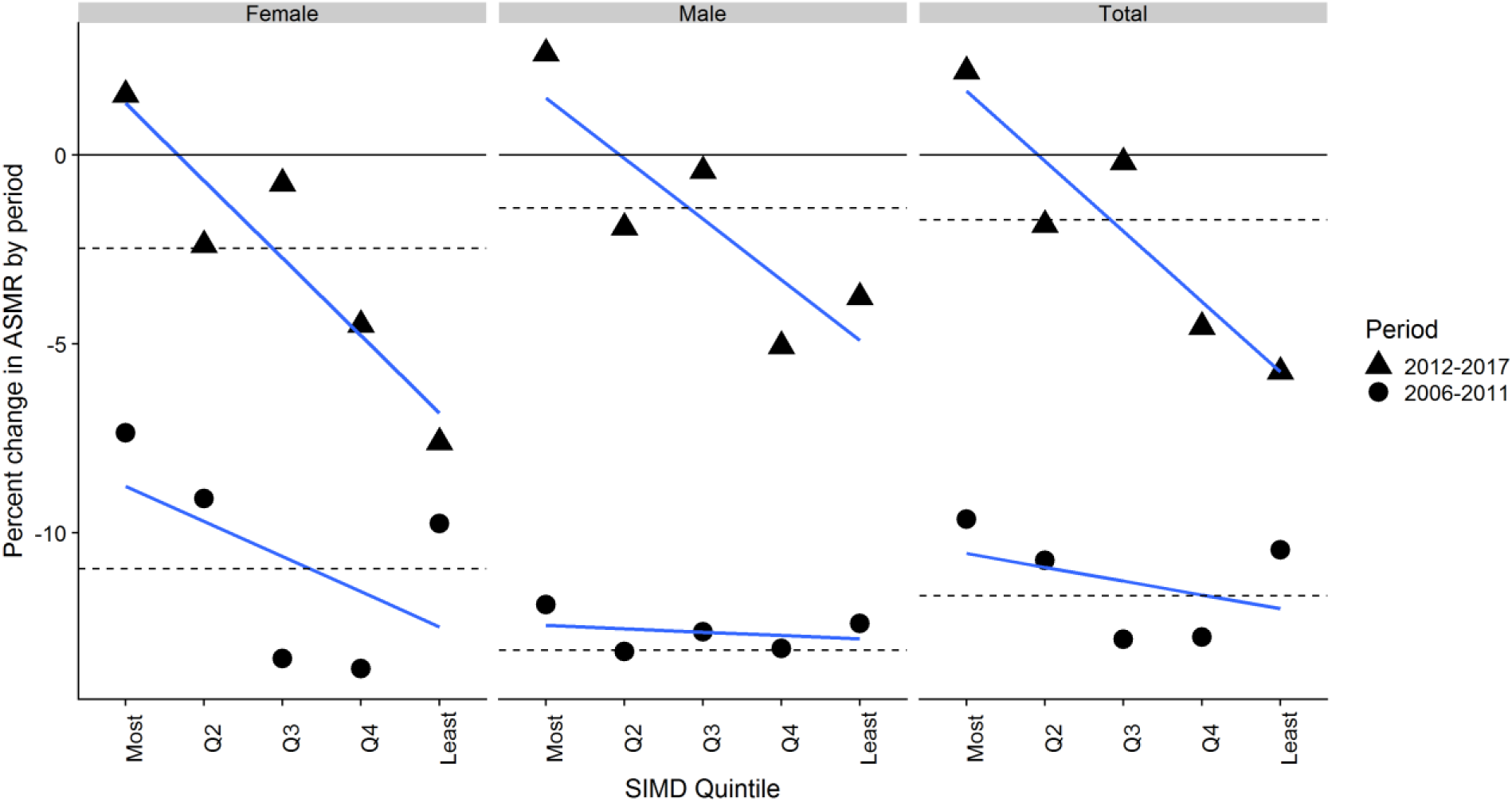
Percentage change in ASMR by period (2006-2010; 2012-2017) by gender and SIMD quintile. Horizontal dashed lines indicate average change in Scotland by period and gender, using fitted rather than observed values.

Table 2 shows the results presented in Table 1, but using the fitted rather than observed values. Although there are some differences, the overall conclusions that may be drawn are the same.

**Table 2.** As with table 1, but using fitted rather than observed values.

Percent change in ASMR by gender, SIMD quintile, and period. (Alternative method)
| gender | period | Percentages |  |  |  |  |  | Model results |  |  |
| --- | --- | --- | --- | --- | --- | --- | --- | --- | --- | --- |
|  |  | Most | Q2 | Q3 | Q4 | Least | Overall | R. sq. | gradient | intercept |
| Female | 2006-2011 | -7.35 | -9.09 | -13.32 | -13.59 | -9.76 | -10.94 | 0.29 | -0.93 (-2.58, 0.72) | -8.76 (-12.80, -4.72) |
| Female | 2012-2017 | 1.60 | -2.38 | -0.75 | -4.50 | -7.60 | -2.47 | 0.85 | -2.05 (-3.03, -1.07) | 1.38 (-1.03, 3.78) |
| Male | 2006-2011 | -11.90 | -13.14 | -12.62 | -13.06 | -12.40 | -13.11 | 0.08 | -0.09 (-0.44, 0.26) | -12.44 (-13.29, -11.59) |
| Male | 2012-2017 | 2.69 | -1.92 | -0.42 | -5.06 | -3.76 | -1.40 | 0.71 | -1.60 (-2.78, -0.43) | 1.51 (-1.36, 4.39) |
| Total | 2006-2011 | -9.63 | -10.73 | -12.82 | -12.76 | -10.45 | -11.67 | 0.16 | -0.37 (-1.31, 0.58) | -10.55 (-12.85, -8.24) |
| Total | 2012-2017 | 2.22 | -1.85 | -0.19 | -4.54 | -5.73 | -1.72 | 0.83 | -1.86 (-2.80, -0.92) | 1.70 (-0.60, 4.01) |

Note:
Overall: Whole of Scotland. R.Sq.: R-Squared for model. Gradient: Increase in % change per unit increase in quintile. Intercept: Predicted % change in most deprived quintile. For gradient and intercept, values in parentheses show lower and upper 95% confidence intervals of coefficients respectively.

### Discussion

This conclusion has shown the effects of using an alternative strategy for calculating ASMR change in two periods, using fitted rather than observed values for the first and last year of each period. Although specific values are not identical, the overall conclusion that can be drawn from the analyses is the same.

## Footnotes

* SIMD quintiles were assigned according to the version of SIMD most relevant to the year in question. Years 2001-2003 use SIMD04, 2004-2006 use SIMD06, 2007-2009 use SIMD09, 2010-2013 use SIMD12 and 2014 onwards use SIMD16.

